# Selective ancestral sorting and *de novo* evolution in the agricultural invasion of *Amaranthus tuberculatus*

**DOI:** 10.1101/2021.07.26.453853

**Authors:** J.M. Kreiner, Amalia Caballero, S.I. Wright, J.R. Stinchcombe

## Abstract

The relative role of hybridization, *de novo* evolution, and standing variation in weed adaptation to agricultural environments is largely unknown. In *Amaranthus tuberculatus*, a widespread North American agricultural weed, adaptation is likely influenced by recent secondary contact and admixture of two previously isolated subspecies. We characterized the extent of adaptation and phenotypic differentiation accompanying the spread of *A. tuberculatus* into agricultural environments and the contribution of subspecies divergence. We generated phenotypic and whole-genome sequence data from a manipulative common garden experiment, using paired samples from natural and agricultural populations. We found strong latitudinal, longitudinal, and sex differentiation in phenotypes, and subtle differences among agricultural and natural environments that were further resolved with ancestry inference. The transition into agricultural environments has favoured southwestern var. *rudis* ancestry that leads to higher biomass and environment-specific phenotypes: increased biomass and earlier flowering under reduced water availability, and reduced plasticity in fitness-related traits. We also detected *de novo* adaptation to agricultural habitats independent of ancestry effects, including marginally higher biomass and later flowering in agricultural populations, and a time to germination home advantage. Therefore, the invasion of *A. tuberculatus* into agricultural environments has drawn on adaptive variation across multiple timescales—through both preadaptation via the preferential sorting of var. *rudis* ancestry and *de novo* local adaptation.

## Introduction

While selection from herbicides is one of the most dramatic and novel selection pressures that new agricultural weed populations experience, a much broader suite of ecological shifts and adaptive changes is likely to accompany the transition into agronomic environments (Murphy & Lemerle, 2006) and resulting range expansion (Clements & Ditommaso, 2011). Weeds that have successfully invaded contemporary landscapes, including crop fields and range lands, are subject to predictable and repeated disturbances, regimented irrigation, extreme interspecific competition, and intensified chemical inputs— all of which should lead to novel selection pressures on life history characteristics (Baker, 1974; De Wet & Harlan, 1975; Vigueira *et al*., 2013). In cereals, weedy ecotypes show greater seedling vigour, loss of germination inhibitors, more determinate growth, and loss of seed shattering in response to selection from harvesting regimes and increased competition (Harlan *et al*., 1973). Weedy populations of *Brassica* have evolved faster flowering outside of their native range (Charbonneau *et al*., 2018) consistent with disturbance regimes selecting for accelerated life history (De Wet & Harlan, 1975; Barrett, 1983; Warwick & Stewart, 2005). Baker (1974) hypothesized many of such traits would be present in an “ideal weed” but additionally, that agricultural regimes may favour phenotypically plastic “jacks of all trades” (Richards *et al*., 2006). With studies of agricultural weeds relatively neglected in evolutionary genetics studies compared to crops despite their important impacts on food productivity (Stewart *et al*., 2009; Ravet *et al*., 2018; Martin *et al*., 2019), the role of plasticity compared to local adaptation, and the relevant timescales of weed evolution to agriculture remains unresolved (Baucom & Holt, 2009). Here, we evaluate the phenotypic and ancestry-dependent basis of agricultural adaptation in native range of common waterhemp (*Amaranthus tuberculatus*), which has experienced widespread shifts in selective regimes along with the contemporary rise of agricultural landscapes.

In addition to *de novo* local adaptation and plasticity, the role of gene flow—especially introgression between wild and domesticated relatives (De Wet & Harlan, 1975)—has been well-recognized in agricultural weed evolution. Hybrid origins of invasive weed populations have been well-documented in the genus *Helianthus* (Kane & Rieseberg, 2008; Muller *et al*., 2011; Lai *et al*., 2012), with multiple wild to weedy transitions occurring via hybridization. In wild and cultivated beets (*Beta vulgaris*), hybridization has led to invasive weed populations with a mix of agriculturally fit traits of both types, including self-fertilization (from the domesticated type), early bolting, and annual flowering (wild type traits) (Arnaud *et al*., 2010). In the present species, common waterhemp, unidirectional gene flow from domesticated *A. hybridus* was identified in a few invasive agricultural weed populations in Illinois, USA (Trucco *et al*., 2009). Beyond the increase in genetic variation driven by introgression from domesticated species—and thus greater opportunity for a response to selection (Fisher, 1930)—the formation of ecotypes is widespread in many common weeds (Brown & Marshall, 1981; Barrett, 1982) and may act as reservoirs of adaptive genetic variation (Baker, 1974).

*Amaranthus tuberculatus* is a diploid annual native to North America (Costea *et al*., 2005). Its tendency to be highly successful in agricultural systems is hypothesized in part result from a combination of its obligately outcrossing dioecious wind-pollinated mating system (Costea *et al*., 2005), extremely high seed production (with females producing on average between 35,000 -1,200,000 notably small (1mm) seeds (Stevens, 1932; Sellers *et al*., 2003; Robert G. Hartzler *et al*., 2004)), and resultantly high levels of diversity (Kreiner *et al*., 2019). Recent inference highlights a massive recent expansion in effective population size over the last century—a key consequence of which is highly parallel target-site resistance evolution (Kreiner *et al*., 2021). In *A. tuberculatus*, two ecotypes have been described, the classification of which has been revised and debated (Riddell, 1835; Sauer, 1955) from one single species (Uline & Bray, 1895; Costea & Tardif, 2003), to most recently, two distinct subspecies on the basis of continuous, clinal morphological variation across their sympatric ranges (Costea *et al*., 2005). We will refer to these lineages as subspecies throughout this manuscript.

The two *A. tuberculatus* subspecies differ in their historical ranges as inferred from herbarium specimens, with var. *tuberculatus* being found along northeastern Missouri and Mississippi water basins, but var. *rudis* (initially circumscribed as *A. tamariscinus*) historically restricted to ruderal habitats in four southwestern states in the USA (Sauer, 1957). The secondary contact of these subspecies over the last two centuries was thought to be driven predominantly by the expansion of var. *rudis* northeastwards. Sauer (1957) hypothesized that the hybridization resulting from this secondary contact led to the agriculturally competitive form. However, he also hypothesized that the hygrophytic nature of species in the genus, and their conditioning to naturally and frequently disturbed riparian habitats, pre-adapted them to the human-mediated disturbances widespread in agricultural landscapes. Preadaptation in the invasion genetics context (*sensu* Liebman *et al*., 2001) refers to prior adaptation in the native range leading to high fitness in novel habitats and is distinct from exaptation which requires a change in trait function (Waselkov & Olsen, 2014; Waselkov *et al*., 2020). Recent genetic and genomic evidence of a longitudinal cline in ancestry between their ancestral ranges supports the secondary contact of *A. tuberculatus* subspecies, but the tendency to see var. *rudis* ancestry in agricultural environments (Waselkov & Olsen, 2014; Kreiner *et al*., 2019) suggests that var. *rudis* in particular, may be pre-adapted. Genome-wide SNP data has been an important tool in understanding the demographic history of the species and population structure across the range, but can further be utilized in cases such as this to better understand features that have facilitated successful establishment and persistence in novel habitats (e.g. Wang *et al*., 2017).

While differences in ecological pressures across natural and agricultural habitats may shape patterns of phenotypic and genomic diversity, this fine-scale evolution is likely to be mediated by geographic gradients in abiotic factors that determine seasonality across broader scales. Adaptive geographic clines in plant traits, latitudinal clines in particular, are ubiquitous and have been widely described across systems for reproductive, defense, and growth related phenotypes (Neuffer, 1990; Stinchcombe *et al*., 2004; Samis *et al*., 2012; Peterson *et al*., 2016; Cornille *et al*., 2018; Bilinski *et al*., 2018; Frachon *et al*., 2018; Exposito-Alonso, 2020). In short day plants (i.e., those where flowering is induced by the shortening of days at the end of the growing season), individuals at higher latitudes should be selected to flower earlier to set seed before frost-induced mortality (Holm, 2010). Longitudinal clines are less common but have been described for flowering time in *Arabidopsis* where it is thought to be associated with geographic variation in winter temperature and precipitation (Samis *et al*., 2008, 2012). Given the latitudinal and longitudinal clines described in *A. tuberculatus* subspecies ancestry, adaptation to these climate gradients may be in part confounded with historical patterns of subspecies divergence.

Here, we test key hypotheses about the role of subspecies hybridization, *de novo* and ancestral variation in facilitating the recent invasion of *A. tuberculatus* into agricultural environments. A recent study performed a replicated common garden experiment using a broad collection of *A. tuberculatus* to test hypotheses about agricultural adaptation, but was largely unable to uncouple geographic and fine-scale environmental drivers of phenotypic differentiation (Waselkov *et al*., 2020). To ensure sufficient power to disentangle broad geographic and environmental drivers of adaptation, we used a paired collection design (Lotterhos & Whitlock, 2015), sampling *A. tuberculatus* in pairs of natural and agricultural sites that were < 25km apart, in a replicated fashion across 3 degrees of latitude and 12 degrees of longitude. We then tested for local adaptation to agricultural environments in a common garden experiment with treatments simulating components of natural and agricultural environments. We performed a water-supplemented treatment and a soybean (*Glycine max*) competition treatment, to simulate riparian and agricultural environments, respectively, along with a control treatment that lacked both competition and water-supplementation. Across collections from 17 sets of paired natural and agricultural populations (34 populations in total), we grew 10 replicates of full siblings from 200 maternal lines across each treatment, totaling to 6000 individuals. Key to testing historical hypotheses about pre-adaptation, we also collected whole genome sequence data from 187 maternal lines to explicitly examine the extent to which subspecies ancestry drives phenotypic differentiation across natural and agricultural environments and geographic clines. By combining a highly replicated phenotypic catalogue and accompanying genomic data, our results provide robust insight into the impact of human-mediated disturbances on trait differentiation and the timescale underlying adaptation to contemporary agricultural environments.

## METHODS

### Collections & P1 Crosses

We made collections of 17 paired populations (a natural and agricultural population collected < 25km apart) in October 2018, from Ohio to Kansas, aiming for 20 maternal lines per population (**Figure 1**). Seed was partitioned into mesh jewelry bags and buried in moist sand for 6 weeks before being grown out, as per stratification recommendations for the species (Leon *et al*., 2007). Four replicates of 700 maternal lines across these populations were sown and grown in growth chambers, under short day conditions to shorten generation time, and germinated under 12 degree temperature amplitude to maximize germination (Leon *et al*., 2004) (16 hours at 32 degrees, 8 hours at 20 degrees). Upon formation of reproductive organs, females and males were immediately bagged to prevent cross-pollination (inspired by McGoey *et al*., 2017) until enough individuals had flowered that controlled, within-population crosses could be conducted. We conducted 345 within-population crosses, where we randomly assigned males to be transplanted into a pot of a female from a different maternal line within the same population, such that we maximized the number of crosses within populations while only performing one cross per maternal line. Upon transplanting the male into the female pot, we bagged the entire above ground portion of the pot and agitated the bags to facilitate male pollen dehiscence. Seeds successfully set in 326/345 crosses, and were harvested for cold treatment prior to the common garden experiment.

**Figure 1.**
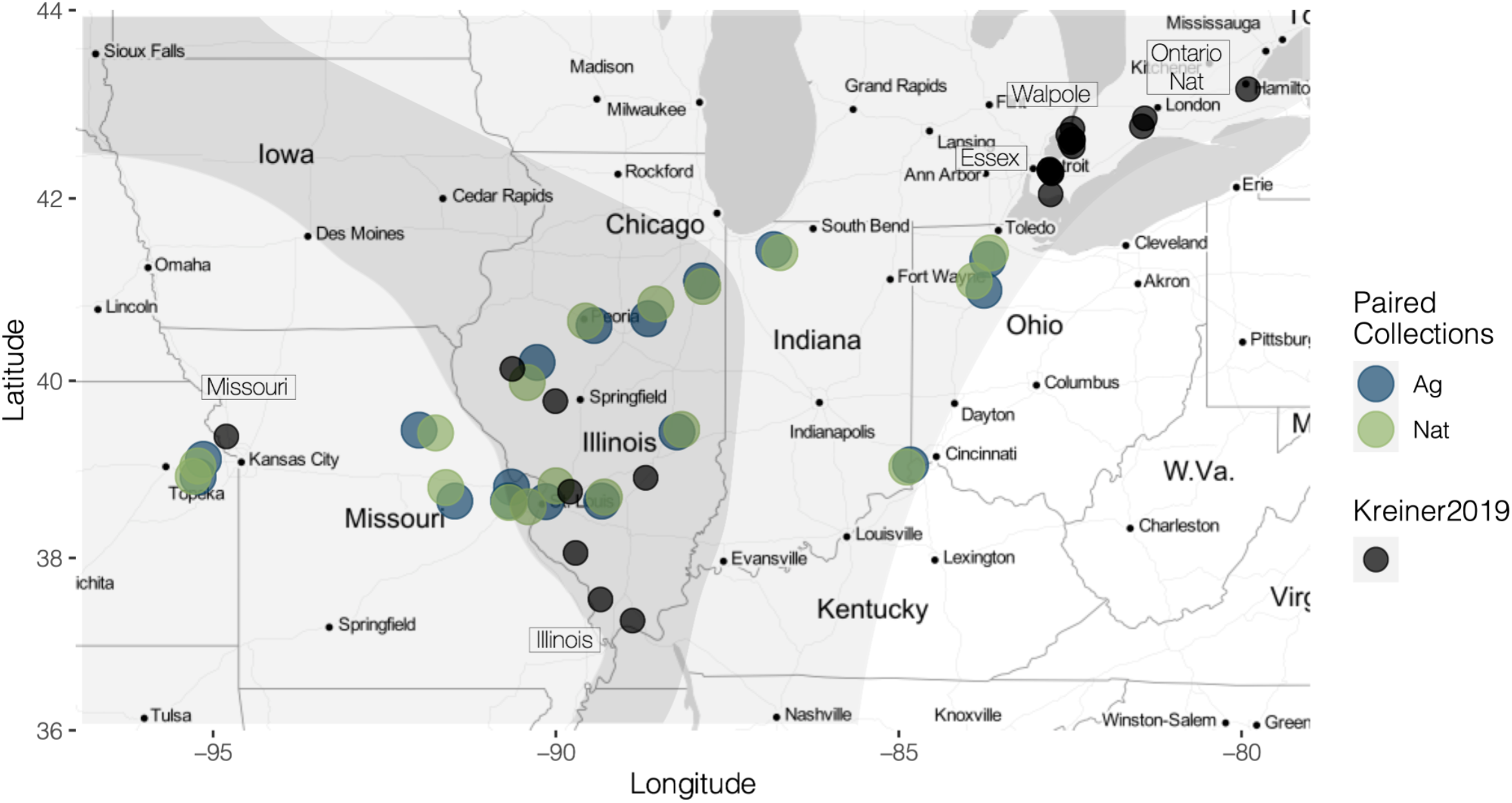
Pairwise collections of natural and agricultural populations spanning the historical sympatric (dark shaded area) and allopatric ranges of *A. tuberculatus* var. *tuberculatus* (northeast) and *A. tuberculatus* var. *rudis* (southwest; range limits adapted from (Sauer, 1957)). For context, we also depict populations along with their regional label from our Kreiner et al., 2019 work in black.

### Roof-top common garden experiment

We subsampled 200 of the 326 F1 lines from within population crosses, with the aim of matching sample size across natural and agricultural environments within each population pair. We cold-treated these lines in a 4°C growth chamber in the dark for 8 weeks, in 8cm wide petri dishes with 7.5ml of deionized water. We sowed seeds in 1L treepots (Stuewe and Sons., Inc) in the greenhouse (July 3rd, 2019), and initially grew them under fluctuating temperatures of 14 degree Celsius at night (8 hours) and 32 degree Celsius in the day (14 hours) to maximize germination. Soybean seed (*Glycine max* var. *dekalb* -DKB, 12-57) was sown in competition pots the next day, with A*maranthus* and soy equally spaced within the pot. Treepots assigned to the water treatment had their bottoms duct taped off, to increase water retention. Plants were watered and checked for germination daily for 10 days, and then moved outside on to the roof of the Earth Sciences Building, Toronto, Ontario, into a fully randomized, complete block design, with every block (10) serving as a replicate of each maternal line (200) in every treatment (x3) (Moved outside July 12th, 2019).

### Phenotyping & data collection

After noting time until germination, we commenced phenotypic measurements at the four-leaf stage starting with cotyledon width (mm), hypocotyl length (mm), and leaf number (in part to control for variation in date of measure). Once the first individual was found in flower, we checked all plants for the start of flowering Monday, Wednesday, and Friday for four weeks. Upon flowering, we also measured stem width, plant height, number of nodes, whether an individual was recorded late (extended inflorescence), or whether the plant had been damaged (these individuals were subsequently excluded from the statistical analyses). Due to a long tail of flowering, after 4 weeks we halved census efforts, alternately checking half of the blocks each Monday, Wednesday, and Friday. Above ground biomass for all undamaged plants was harvested into paper bags, starting 8 weeks after the start of flower and lasting until 11 weeks after flowering until all plants had been harvested. Upon harvest, we recorded sex, flower color, and stem color. Plants were then dried in a 50°C oven for 3 days and weighed for above ground biomass. In total, we measured 11 quantitative phenotypes: days until germination, cotyledon width, hypocotyl height, time to flowering, height at flowering, node number at flowering, stem width at flowering, sex, flower color (visual rating on a scale of 1 [light green] to 4 [dark purple]), stem color (visual rating on a scale of 1 [light green] to 4 [dark purple]), and dry biomass. We also tracked greenhouse number, greenhouse block, roof block, days to measurement, and days to harvest.

### DNA collections & Sequencing

We sampled 2-3 of the youngest leaves on each individual in two blocks of our common garden experiment just before harvest. Leaves were immediately put in tubes and submerged in liquid nitrogen before being stored at -80 °C until extraction. We extracted DNA from the 200 unique maternal lines that were grown in the common garden experiment. Total DNA was extracted using Qiagen DNeasy plant mini kit according to manufacturer’s instructions. We sent DNA samples to Genome Quebec Innovation Centre (McGill University), Montréal, QC, Canada for library preparation and sequencing; 187 ended up being sequenced due to extraction and library quality. Libraries were prepared using the NEB Ultra II Shotgun gDNAlibrary preparation method and sequenced on four lanes of Illumina NovaSeq S4 PE150 (2 × 150) sequencing platform using 96 barcodes. A total of ∼25 billion reads (25,818,840,892) were generated, with an average of ∼137 million (137,334,200) per individual.

### Mapping & SNP calling

We aligned reads to the female *A. tuberculatus* reference genome (Kreiner *et al*., 2019), using BWA-mem v0.7.17-r1188 (Li, 2013). After mapping, individuals had an average diploid coverage of 28X. Duplicate reads were removed with picard MarkDuplicates (Broad-Institute, 2016). We used freebayes v1.1.0-46 (Garrison et al., 2010) to call SNPs using default settings except for --max-complex-gap 1, --haplotype-length 1, and --report-monomorphic. We then filtered SNPs such that sites were removed based on excess missing data (>20%), dustmasked for low complexity, removed multiallelic snps and indels, allelic bias (AB <0.25 and >0.75), and on overall variant call quality (QUAL < 30, removing sites with greater than a 1/1,000 genotyping error rate). Furthermore, since high coverage data tends to overestimate mapping quality (such that it no longer scales with depth), we followed recommendations in (Fang, 2014), further removing sites with a particularly high depth (mean depth + 3(sqrt(mean))), that have a qual score < 2*depth. Five genotypes were removed from downstream analyses due to >5% sequencing error rate based on a KMER based analysis (Ranallo-Benavidez *et al*., 2020), resulting in a total of 20,555,154 SNPs.

### Population Structure

We merged filtered, high quality SNPs from the 182 high quality resequenced genomes, with the high quality SNP set from (Kreiner *et al*., 2019). Briefly, these previous collections were made from 8 agricultural populations in Illinois and Kansas with reports of high levels of resistance to herbicides from farmers, and fields within Walpole Island and Essex County, Ontario, Canada, where new reports of problematic agriculturally associated populations of *A. tuberculatus* have only recently occurred in the last decade. Additionally, this dataset included 10 individuals collected from nearby Ontario natural populations, occurring alongside the Thames River outside of London, and the Grand River outside of Hamilton. From this merged set of 2,591,759 SNPs, we investigated population structure with a principal component analysis in plink (option --pca) (Purcell *et al*., 2007). We investigated predictors of genome-wide relatedness in a multiple regression framework using PC1 and PC2 as dependent variables and longitude, latitude, sex, environment, and population pair as independent variables. To test if predictors were different among PCs, we used a grouped regression approach, testing whether the value of the principal component was predicted not just by these same predictors but the interaction of these predictors and principal component number (i.e. PC1 or PC2). Lastly, we used the program Faststructure (Raj *et al*., 2014) to estimate the proportion of individual and population admixture levels, at K=2 (testing *a priori* hypotheses about the distribution of subspecies variation) and for comparison, at K=3.

### Linear mixed models and tests for preadaptation

#### Modelling individual level phenotypic variation

We used R to estimate linear mixed models (implemented with the package lme4) of geographic, environmental, and sex-based predictors of each of our 11 phenotypes measured in the common garden experiment, all of which were evaluated with a type III sums of squares. For analysis of phenotypic variation at the individual level from the common garden experiment, we accounted for the relevant block effect (typically roof block, except for time to germination, for which we used greenhouse: greenhouse block), and nested hierarchical structure of maternal lines (family) within populations as random effects. For all 11 traits, individual level phenotypic variation was measured with the following model structure (note that we initially included an environment by treatment interaction but removed it due to low explanatory power and lack of significance for all phenotypes):

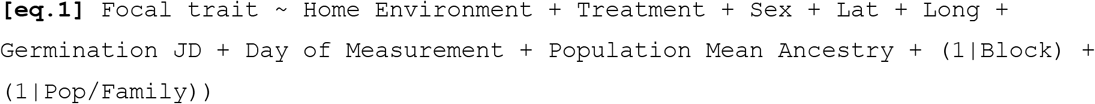

Our fixed effect predictors had the following characteristics: home environment had two levels (natural or agricultural), treatment had three levels (control, soy, or water), sex had two levels (M/F), and latitude and longitude were both treated as continuous variables, referring to the geographic coordinate of the originating population. Except for when we were modelling germination itself, Julian day of germination and Julian day of measurement were included as covariates to account for variation in how long a plant had been growing prior to measurement. We used the population-mean subspecies ancestry (the average of the faststructure inferred proportion of an individual’s genome assigned to cluster 1 at K=2 across all sequenced individuals within a population) as an estimate of genetic structure in these models. We used population level estimates rather than family level because of low sequencing replication (1 individual per full sibling family in an obligately outbreeding species), and because population level estimates should reflect broad scale geographic patterns in ancestry, similar to using latitude and longitude as proxies for geographic and climatic variation. We compared the full model as shown above to a reduced model that excluded population mean ancestry using AIC to evaluate its importance in explaining phenotypic differentiation.

#### Testing the role of var. rudis ancestry

We were interested in explicitly testing *a priori* hypotheses about the role of var. *rudis* ancestry on adaptive phenotypic variation, given the work of Sauer (1957) and Waselkov (2014). To incorporate both ancestry and phenotypic traits, we analyzed the sex-specific phenotypic means within each treatment, within each maternal line. Thus while phenotypic family means were distinct across each sex and treatment level with a maternal line, we assigned all treatment and sex replicates of a maternal line the Faststructure ancestry estimate we attained from their single sequenced full-sibling. We then examined two key fitness related traits, biomass and flowering time, separately for males and females given strong sexual dimorphism in the species.

Beyond the direct linear effect that var. *rudis* ancestry might have on phenotypic variation, we were particularly interested in testing whether there was a positive quadratic effect on fitness-related variation (indicating a role for heterosis), and whether there was an interaction between the proportion of var. *rudis* ancestry and experimental treatment on fitness-related variation (indicating a role for pre-adaptation, in that ancestry confers environment-specific benefits). Lastly, we wanted to test the extent to which phenotypic variation differed among environments, regardless of ancestry (indicating *de novo* agricultural adaptation). To test these three alternative but not mutually exclusive hypotheses, we used a model similar to one testing linear selection gradients separately for males and females, regressing these ancestry and environment terms along side standardized phenotypic predictors on two fitness related traits (flowering time and biomass) to account for correlated trait evolution:

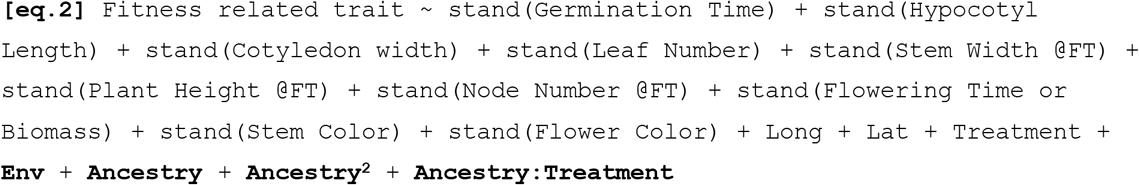

## RESULTS

### Drivers of population structure and subspecies-specific variation

To understand patterns of genetic relatedness that underlie phenotypic variation within and between populations, and the potential role of subspecies ancestry in facilitating agricultural adaptation, we first characterized patterns of population structure and ancestry across our accessions in the context of previously characterized populations (**Figure 1**).

We find that individuals from our paired-environment collections show a longitudinal cline in ancestry, as expected (Sauer, 1957; Waselkov & Olsen, 2014; Kreiner *et al*., 2019) **(Figure 2A,C)**. Our previous work showed that Ontario Natural populations in the eastern part of the range are homogenous for var. *tuberculatus* ancestry, and along with our most westerly collections in Missouri, that nearby Essex county agricultural populations are homogenous for var. *rudis* ancestry, likely reflecting a long-distance introduction event from the Midwest (Kreiner *et al*., 2019). The nearly 200 genotypes we have added to this genome-wide inference of population structure supports the circumscription of the historical ranges of these two subspecies, in that northeastern populations (e.g., Maume, Mccombe, Weston) showed a higher proportion of *Amaranthus* var. *tuberculatus* ancestry while southwestern populations showed predominantly var. *rudis* ancestry **(Figure 2C)**. A joint PCA of common garden accessions and samples previously characterized in (Kreiner *et al*., 2019), illustrates that common garden accessions showed somewhat less extreme population structure along PC1 and PC2 of genome-wide allele frequencies, consistent with the more continuous but geographically intermediate sampling we performed **(Figure 2A)**. Individuals from our common garden typically fell in a very similar position for PC2 and showed much more variation along PC1, which explained 18% of the total variation in genotype composition across the 349 joint accessions. We have previously shown PC1 to strongly reflect subspecies ancestry (Kreiner *et al*., 2019).

**Figure 2.**
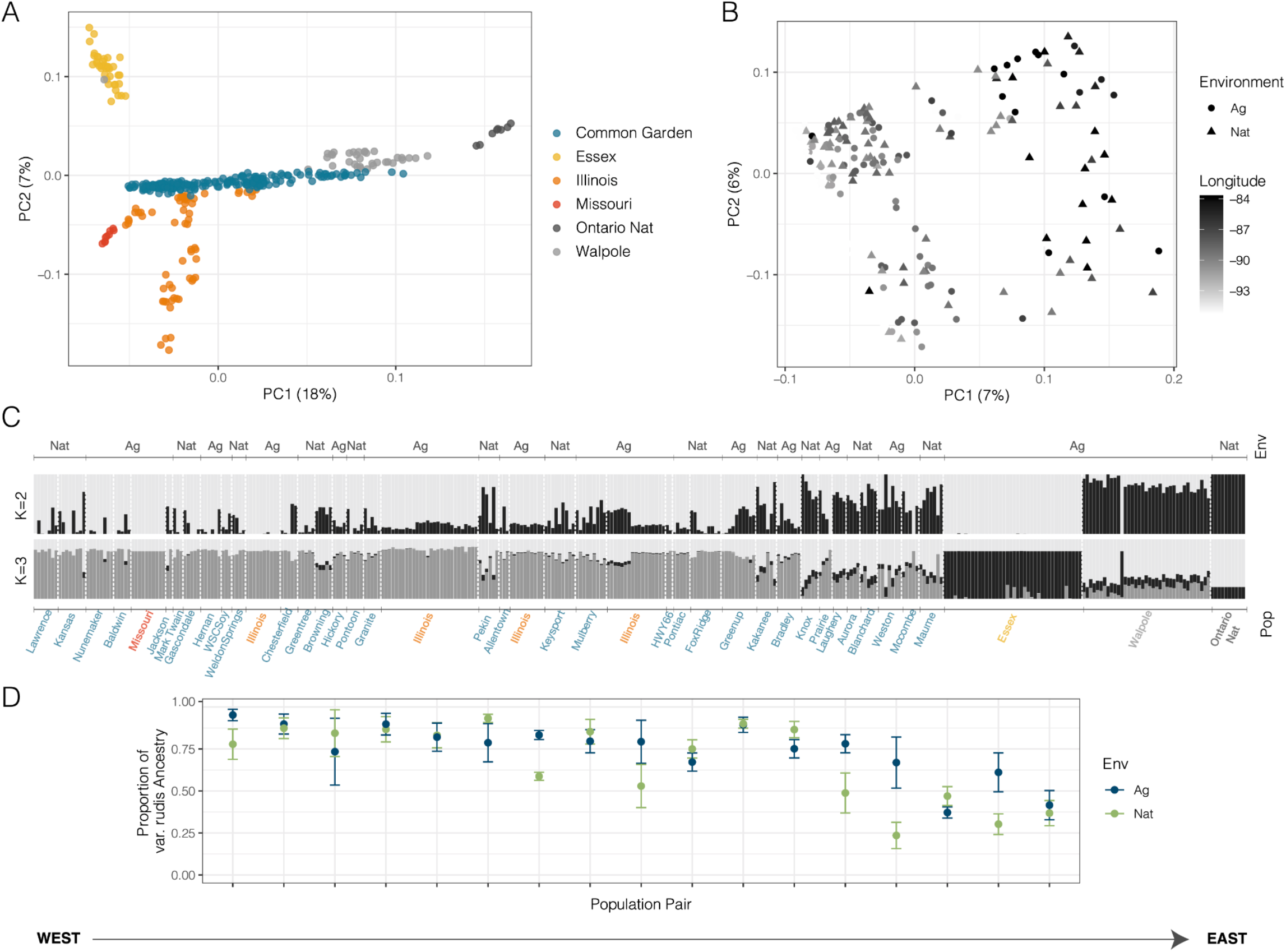
Patterns of population structure from pairwise collections, grown in the common garden experiment, in the context of broader sampling from. (Kreiner *et al*., 2019). **A**) Principal component analysis of samples from (Kreiner *et al*., 2019), where populations have been identified as homogenous for both *A. tuberculatus* var. *tuberculatus* ancestry (e.g., Ontario Nat) and *A. tuberculatus* var. *rudis* ancestry (e.g. Kansas) along with 187 genotypes grown in the common garden experiment originating from pairwise Nat-Ag population sampling. **B**) Principal component analysis of just common garden genotypes. **C**) Population structure (at K=2, reflecting ancestry of var. *rudis* in black and var. *tuberculatus* in grey) across both previously analyzed samples and common garden accessions. Plot is sorted by longitude (from west to east) and labels of populations from Kreiner et al., 2019 are colored according to the legend in A). **D**) Higher proportion of average var. *rudis* ancestry in agricultural versus natural environments in population pairs, sorted by longitude (west to east). Error bars represent standard error.

When we performed a PCA exclusively on genotypes from our common garden experiment, PC1 in comparison explained 7% of the variation in genotype composition **(Figure 2B)**. From a multivariate regression of this PCA of just common garden accessions, we find that PC1 significantly relates to longitude (*F*1,180 = 27.05, *p* < 0.001), population pair (*F*1,180 = 4.58, *p* = 0.03), and environment (agricultural vs. non-agricultural, *F*1,180 = 5.51, *p* = 0.02), but neither latitude nor sex. To test if the predictive effects of environment differ across PCs, we performed a follow up grouped-regression approach jointly examining if the value of the first two PCs can be explained by environment (agricultural vs. natural), pair, and longitude, and whether those predictors interact with PC number (i.e., whether predictive effects differ across PCs). This grouped model fails to detect a significant environment by PC interaction, implying that the predictive effects of environment are consistent across multiple dimensions of allele frequency differentiation, but picks up a significant pair x PC (*F*1,366=10.3686, *p* = 0.001396) and longitude x PC interaction (*F*1,366=84.95 p = < 2.2e-16) with both pair and longitude better predicting the first PC. After accounting for these interactions, we find that the linear effects of pair (*F*1,366=9.37, *p* = 0.0024), longitude (*F*1,366=94.7353, *p* = < 2.2e-16), and environment (*F*1,366=3.90, *p* = 0.049) are significant predictors of values of the first two PCs of genotype composition.

The influence on environment (natural or agricultural) on ancestry identified in the common garden specific PCA is apparent in **Figure 2C**, where agricultural populations show an excess of var. *rudis* ancestry given their longitude, and more apparently so in **Figure 2D**, within their population pair—a more direct comparison of environmentally-driven sorting of ancestry. Indeed, a multivariate regression of the Faststructure inferred proportion of var. *rudis* ancestry (grouping 1 at K=2) finds longitude (*F*1,167=8.29, *p* = 0.005), pair (*F*1,167=3.25, *p* = 6.70e-05), and environment (F1,167=6.66, p = 0.011) to be significant predictors of ancestry, with more var. *rudis* ancestry in agricultural environments. On average, this pattern resulted in a 7.8% excess of var. *rudis* ancestry in agricultural environments across all population pairs, after controlling for other covariates. Of our 17 population pairs, 9 show greater var. *rudis* ancestry in agricultural environments, with the median proportion of var. *rudis* ancestry being 25% greater in these agricultural populations (and up to 43% greater in the most extreme pairing; **Figure 2D)**. In contrast, the remaining 8 pairs that have greater var. *rudis* ancestry in natural environments differs by only 8% on average (and at 14% at its maximum). The significant enrichment of var. *rudis* ancestry given a population pair’s longitude (see PC1 regression) supports the hypothesis that the expansion of the var. *rudis* contributed to the *A. tuberculatus* agricultural invasion (Waselkov & Olsen, 2014; Waselkov *et al*., 2020). Furthermore, that population pair significantly predicts ancestry across a disparate sampling suggests that selection is maintaining this pattern of environment-dependent ancestry despite nearby natural and agricultural populations being highly connected through gene flow.

### Plasticity, sexual dimorphism, and genomic trait variation

#### General observations from the common garden experiment

Across the 4493 individuals fully phenotyped in the common garden experiment, we found almost a perfect 1:1 sex ratio (2252 males vs. 2241 females). On average, we completely phenotyped 22.5 maternal lines per population with 11.2 females per maternal line (sd = 3.17) and 11.26 males per maternal line (sd = 3.33), with an average of 7.5 maternal line replicates phenotyped across each of three treatments.

#### Reaction norms & treatment effects

Our treatments worked as expected, with population-mean flowering time and dry-biomass reflecting that plants generally grew larger and flowered fastest in the water treatment, and were smallest and later flowering in the soy competition treatment **(Figure 3)**. However, rather than recapitulating the often-flooded environment of natural populations, our water treatment only tended to reduce drought stress relative to the control treatment. As an example of the magnitude of the effect of our three treatments, we characterized how flowering time differed depending on whether a genotype was reared in the wet, control, or soy treatment. A least squared mean estimate from the general regression model, using flowering time as a response variable, estimates that the water treatment led to 1 day earlier flowering than the control and 4 days earlier than the soy treatment. We further modelled phenotypic plasticity as a random effect, testing for a family by treatment interaction (lmer notation = 1 treatment:family).

**Figure 3.**
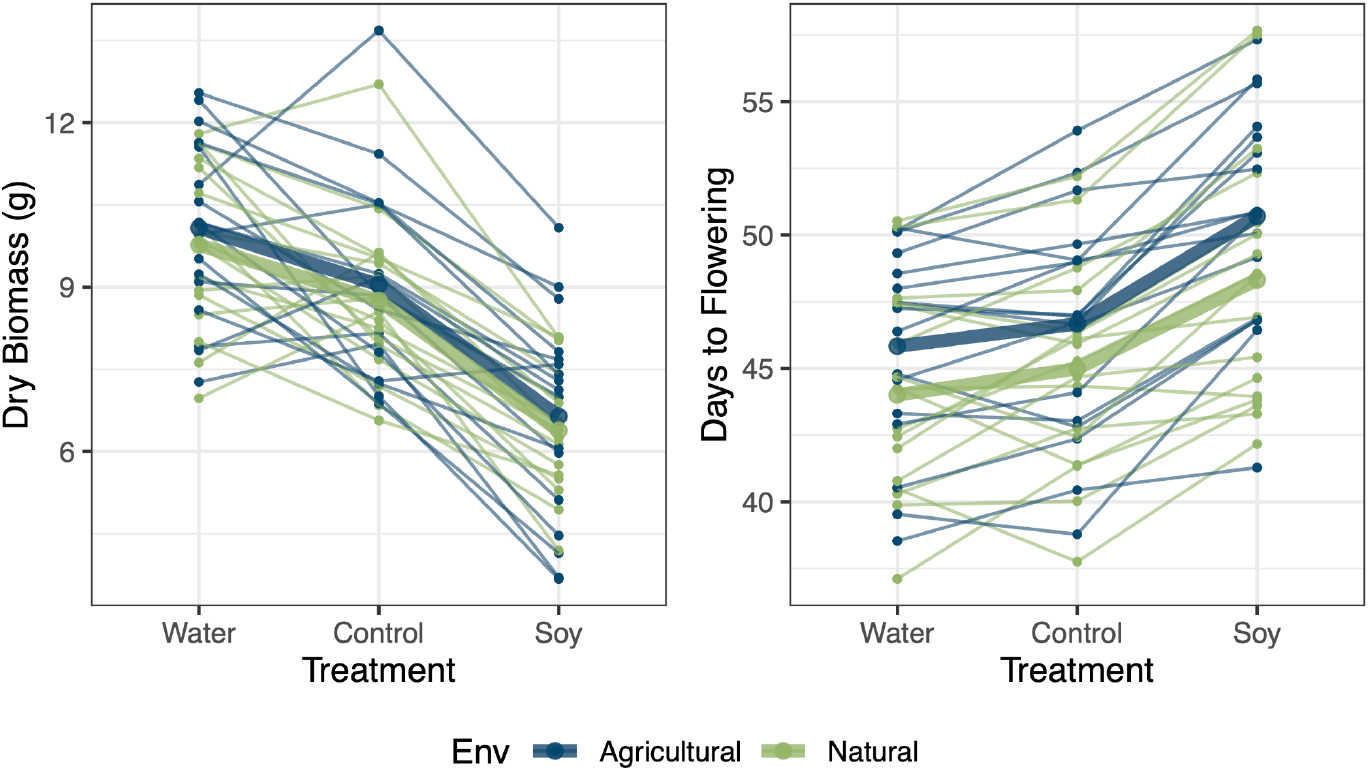
Population-level reaction norms of biomass and flowering time to the water, control, and soy treatment in the common garden experiment. Thin and thick lines represent population and environmental reaction norms, respectively, and are additionally coloured by whether collections were found in natural or agricultural environments.

Following the general model additionally including this plasticity random effect term, we find that modelling phenotypic plasticity results in a significantly better fit to our data despite the additional degrees of freedom (χ^2^_df=1_ =369.27, *p* < 0.001) and can explain an additional 5.28% of the variation in flowering time (on top of the 51% of the base model; conditional r^2^)—implying 10% of the explainable variation in flowering time is plastic. Phenotypic plasticity was of even greater importance for determining dry biomass, explaining an additional 11% of variation in dry biomass on top of the 64% that can be explained in our base model (χ^2^_df=1_ =320.3824, *p* =<2e-16), implying that ∼16% of the explainable variation in above ground biomass in plastic. To test whether populations from natural and agricultural environments differed in the extent of phenotypic plasticity, we compared our plasticity model to one that allowed plasticity to differ across environments (lmer notation = Environment | Treatment:Family). For both biomass and flowering time, allowing plasticity to vary among environments did not increase the variance explained in the model, with environmental differences in plasticity not significantly explaining dry biomass (χ^2^_df=2_ =0.3285, *p* =0.8485) and very marginal effects on flowering time (χ^2^_df=2_ =4.613, *p* =0.0996) (**Figure 3**).

### Drivers of phenotypic variation and the role of subspecies ancestry

#### Geographic, environmental, and subspecies trait divergence

We found evidence for phenotypic differentiation across our broad sampling of *A. tuberculatus* individuals grown in the same common garden, by latitude, longitude, sex, and between agricultural and natural environments. We evaluated phenotypic variation across all individuals in the common garden experiment in the typical manner of controlling for nested family structure, but also considering the effect of accounting for subspecies specific ancestry. For all models, adding population-mean ancestry as a covariate led to a substantially smaller AIC (**Table 1**).

**Table 1.** Summary of the results of linear mixed models for 11 focal traits measured across three life history stages, comparing the full model to a reduced model that drops population-mean ancestry as a covariate. Data were analyzed at the individual level using nested family structure and block effects as random effect covariates. The last column depicts predictors that change in p-value class with the addition of ancestry, and their respective shift in their χ^*2*^ test statistic. Significance codes: *** = p < 0.001, **= p < 0.01, * = p < 0.05, . = p < 0.1

| Early Life History Traits | Significant Predictors | ΔAIC (full-red.) | Predictors that change p-value class with +Ancestry ($\Delta X^2$) |
| --- | --- | --- | --- |
| Days to Germination | Treatment (.), **Env x Treatment** (**), Sex (**), **Anc.** (*) | -407.40 | NA |
| Hypocotyl Length | Treatment (***), Lat (***), **Anc** (**) | -515.50 | NA |
| Cotyledon Width | NA | -299.64 | Environment (2.87→1.57) |
| Early Leaf Number | Treatment (***), Long (.), Lat (.), Sex (**) | -654.74 | Environment (2.86 →1.41), Sex (6.47→7.29) |
| **Mid-life History Traits** |  |  |  |
| Height at Flowering | Treatment (***), Lat (***), Sex (***) | -642.48 | NA |
| Node Number at Flowering | Treatment (***), Long (*), Sex (***) | -484.63 | NA |
| Stem Width at Flowering | Treatment (***), Lat (***), Sex (***), **Anc.** (***) | -330.20 | Latitude (5.39→13.91) |
| Days to Flowering | **Env** (.), Treatment (***), Sex (*), Lat (**), Long (*), Sex*Lat (*) | -574.04 | Longitude (8.07→4.26), Latitude (11.02→10.53), Environment (4.38→3.04) |
| **Late-life History Traits** |  |  |  |
| Flower Color | Treatment (*), Sex (***), **Anc.** (.) | -154.04 | NA |
| Stem Color | Treatment (***), Sex (***), **Anc.** (***) | -207.48 | Longitude (13.52→0.64) |
| Dry Biomass | Treatment (***), Sex (***), Treat x Sex (***), Lat (*), **Anc** (.) | -452.90 | Latitude (3.00→4.64) |

Despite sampling a far greater range of longitude than latitude, after accounting for ancestry, latitude more consistently predicted variation in our measured traits (6/11 traits significantly predicted by latitude, versus 4/11 for longitude: **Table 1**). For longitude, accounting for subspecies-specific ancestry tended to decrease its explanatory power, considerably decreasing the longitude χ^2^ and removing its significant effect in explaining days to flowering and stem colour (**Table 1, Figure 4**). While latitude also covaried with ancestry, accounting for ancestry tended to increase the explanatory power of latitude (e.g., for stem width at flowering and dry biomass) (**Table 1, Figure 4**). The observation that ancestry absorbs more explanatory power of longitude compared to latitude is consistent with the stronger longitude clines in ancestry we describe above.

**Figure 4.**
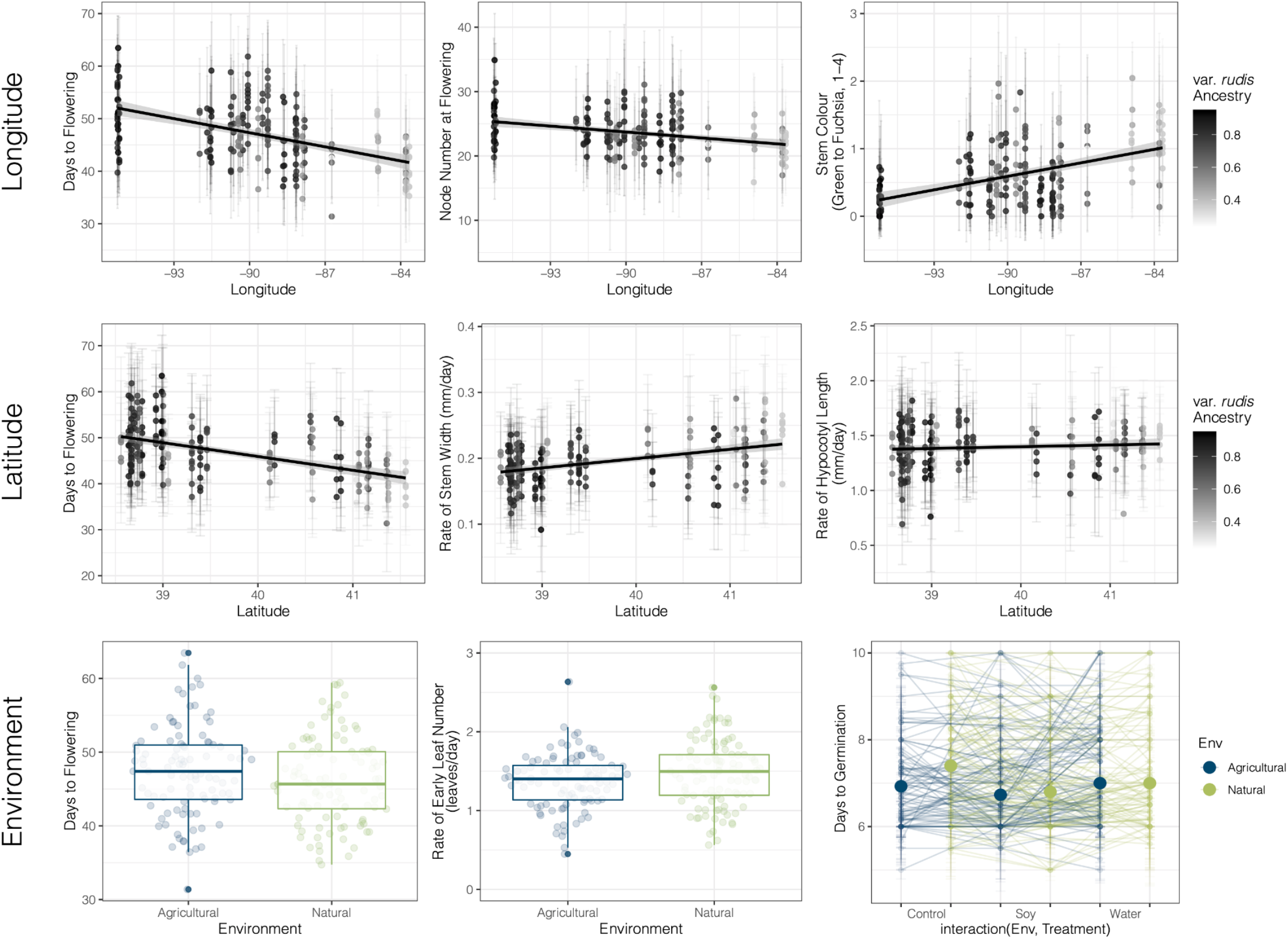
Phenotypic differentiation by longitude, latitude, environment, and the confounding effect of ancestry. All points indicating family-wise means. **Top**) Days to flowering, node number at flowering, and stem color by longitude and proportion of var. *rudis* ancestry. **Middle**) Days to flowering, rate of stem width, and rate of hypocotyl length by latitude and proportion of var. *rudis* ancestry. **Bottom**) Days to flowering and rate of early leaf number (leaves/day) by environment (Left). Germination Julian Day, with large points indicating the mean of the environment by treatment interactions and connecting lines representing family-mean reaction norms (Right). For clarity, the days to germination family-mean reaction norms represent data for males only.

Subspecies ancestry itself significantly predicts days to germination (χ^2^ =4.1204, *p* =0.04237), hypocotyl length (χ^2^ =9.6870, *p* =0.0018558), stem width at flowering (χ^2^ =13.8240, *p* =0.0002008), stem color (χ^2^ =26.0079, *p* =3.400e-07), and marginally flower color (χ^2^ =3.7162, *p* =0.05389) and dry biomass (χ^2^ =3.4895, *p* =0.0617580) in our individual level regressions (**Table 1**)—highlighting the role of the historical isolation between these two lineages in shaping current day patterns of phenotypic variation from early to late life history. We found little signal of the classic reciprocal common garden test for agricultural adaptation across our measured traits—a home environment advantage—however days to germination showed a significant treatment by environment interaction (χ^2^ =9.3759, *p* =0.009), with agricultural types taking the longest to germinate in wet conditions and natural types taking the longest to germinate in control conditions (germination in both environments was shortest in the soy treatment) (**Figure 4**). With genetic ancestry showing significant differences between home environments, we tested the extent that accounting for ancestry would further resolve natural-agricultural phenotypic differentiation. Before accounting for ancestry, we found that agricultural types tended to have marginally wider cotyledons, fewer leaves early on, and a significantly longer time to flowering. After accounting for ancestry, environment only remained a marginally significant predictor for days to flowering (χ^2^ =3.0378, *p* =0.081347) (**Table 1, Figure 4)**. Thus, while we find evidence of agricultural adaptation via germination, fine-scale evolution in agricultural environments additionally tends to be drawing preferentially on standing variation across *A. tuberculatus* subspecies that has been shaped by allopatry and adaptation to geographic gradients.

#### Subspecies ancestry effects on fitness-related traits: tests for pre-adaptation, de-novo adaptation, and hybrid vigour

We hypothesized that if var. *rudis* ancestry is pre-adapted to agricultural habitats, the effect of the proportion of var. *rudis* ancestry on key life history characteristics such as biomass and flowering time would vary depending on experimental treatment. In contrast, if hybridization between subspecies has facilitated much of the *A. tuberculatus*’s contemporary invasion through hybrid vigour, we predicted that var. *rudis* ancestry would have a non-linear relationship with fitness related traits. Finally, if populations were adapting to agricultural regimes *de novo*, we predicted that fitness related traits should vary among natural and agricultural environments, regardless of ancestry.

An analysis at the family-mean level of lifetime above-ground biomass in males found no quadratic effect of var. *rudis* ancestry, however we found a significant linear effect of the proportion of var. *rudis* ancestry, where pure var. *rudis* types were predicted to accumulate biomass at a rate of 0.046 g/day more than pure var. *tuberculatus* types (*F*1,498=3.9112, *p=*0.0485153). Additionally, the interaction effect of var. *rudis* ancestry by treatment and marginally, environment, significantly affected male biomass (Ancestry x Treatment: *F*2,498=3.335, *p=*0.0364; Environment: *F*2,497=3.234, *p=*0.0728). Biomass-based fitness estimates were lower in males from natural environments regardless of treatment (**Figure 5)**. The significant interaction between var. *rudis* ancestry and treatment revealed that the proportion of var. *rudis* ancestry had little effect on biomass in the water and soy treatment, but that it substantially increased biomass in the absence of competition and water supplementation (**Figure 5)**. For male flowering time, of our interest in linear, non-linear, and interaction effects of ancestry, only the interaction between var. *rudis* ancestry and treatment remained a significant predictor (Ancestry x Treatment: *F*2,498=3.2144, *p=*0.041015), with the effect of ancestry on flowering time in soy significantly different from that in the control treatment (*t =* 2.514, *p* = 0.0123). While higher levels of var. *rudis* ancestry led to a shorter time to flowering in the control treatment, increasing var. *rudis* ancestry extended time to flowering in the soy treatment (**Figure 5)**. These male, treatment-specific effects of ancestry more broadly reflect a loss of phenotypic plasticity (convergence of time to flowering or rate of biomass, regardless of reared environment) with increasing proportion of var. *rudis* ancestry. Female flowering time and biomass were not influenced by var. *rudis* ancestry (neither linear or quadratic terms, or through its interaction with treatment), but females showed marginally lower biomass and marginally earlier time to flowering in natural compared to agricultural environments (Biomass: *F*1,498=3.091, *p=*0.0793; Flowering: *F*1,498=3.374, *p=*0.0668).

**Figure 5.**
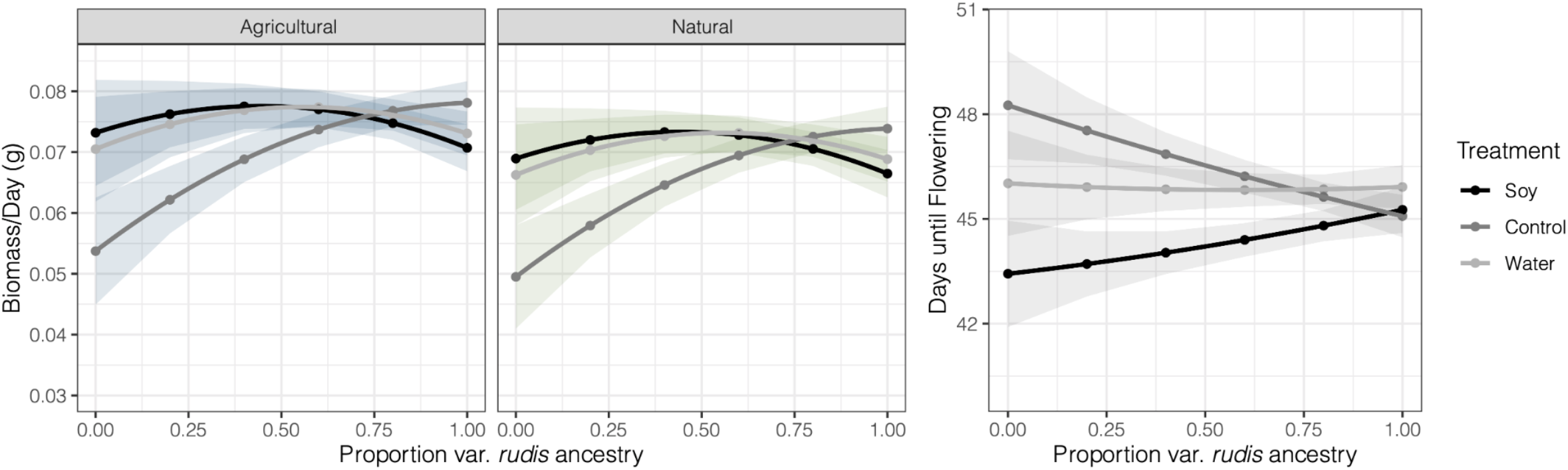
Interaction between proportion var. *rudis* ancestry and experimental treatment predicts male biomass (left) and male days to flowering (right). Results are based on the least squares means of multiple regressions that controlled for the indirect effects of all other measured phenotypes on fitness. The effect of home environment on biomass is apparent in comparing the two biomass panels. Note that the non-linear effect of var. *rudis* ancestry does not significantly predict either biomass or days until flowering.

## DISCUSSION

We found evidence of phenotypic differentiation in *A. tuberculatus* across geographic gradients, across subspecies, and resulting from the transition from natural riparian to highly disturbed agricultural environments. Geographic variation in climate has led to strong latitudinal clines in growth and life history characteristics, however the longitudinal phenotypic clines are in large part explainable by subspecies ancestry. While we found that agricultural populations tended to exhibit shorter times to germination in their simulated “home” environment, slower early growth rates, and longer time to flowering—these differences were also in part related to differential subspecies ancestry across environments. We found that the transition of *A. tuberculatus* into agricultural environments has favoured southwestern var. *rudis* ancestry—ancestry that leads to lower phenotypic plasticity in fitness related traits and generally higher biomass, but also environment dependent phenotypes. Higher var. *rudis* ancestry results in longer time to flowering in the face of competition, and both faster time to flowering and increased biomass in environments lacking competition or water supplementation (i.e. the control treatment). When accounting for these complex treatment-dependent effects of ancestry, we also found marginally lower biomass and earlier flowering time in natural compared to agricultural environments, suggesting invasive agricultural populations may be adapting to a new fitness peak. Therefore, phenotypic differentiation among natural and agricultural environments that are likely to be driven by altered selection regimes have in part been driven by the selective sorting of var. *rudis* ancestry (pre-adaptation), and in part been driven by *de novo* adaptation. These results highlight how human-mediated disturbance and agricultural regimes drive the evolution of native species, shaping interactions between once isolated lineages and drawing from adaptive variation on multiple timescales.

### Phenotypic and genomic underpinnings of agricultural adaptation

We were interested in the extent to which phenotypic variation consistently differed among natural and agricultural environments in a common garden of highly replicated genotypes from environmentally paired populations across a wide sampling of the *A. tuberculatus* native range. An initial investigation of predictors of individual level phenotypic variation observed in our common garden experiment, accounting for hierarchical structure of families within populations, showed that agricultural populations tend to flower later (1.5 days), have a fewer number of leaves early on in their life history, and suggested local adaptation via germination (home advantage, through an environment x treatment effect) of days to germination (**Figure 4**). Agricultural types tended to germinate latest in the water treatment, whereas natural types germinate latest in the control treatment (lacking both competition and water supplementation), suggesting that agricultural adaptation via germination may be driven by moisture availability rather than competition. Interestingly, in large part these phenotypic differences were not consistent with the hypothesis that disturbance regimes will select for accelerated life history in agricultural populations (De Wet & Harlan, 1975). Overall, while the magnitude of these phenotypic differences between environments appears small, that we observed consistent differences across environments with our paired collections and accounting for ancestry suggests that selection may be acting on these traits despite little observed evolutionary response. The efficacy of the evolutionary response to agriculture may be dampened by very recent timescales of selection, or gene flow hindering an environment-specific response to selection, as suggested by population pair significantly predicting patterns of population structure across our collections in regressions of genomic PCs and faststructure predicted ancestry (**Figure 2**).

### A role for both pre-adaptation via preferential sorting of ancestry & de novo agricultural adaptation

Gene flow across environments and across the range may not only lead to reduced differentiation in phenotypes but may also drive heterogeneity in shared ancestry. In the case of *A. tuberculatus* subspecies, secondary contact between these two ancestral lineages has led to longitudinal clines in ancestry across their range. We find that in large part, the longitudinal clines in phenotypes we initially observed (e.g.days to flowering, rate of node number, stem color) covary with longitudinal clines in ancestry (**Figure 2, Figure 4**), implying that differences in the historical ranges of *A. tuberculatus* subspecies has resulted in considerable phenotypic differentiation in not just seedling morphology and seed dehiscence as has been described (e.g. Costea *et al*., 2005), but also in several life history characteristics. Stem color shows the most extreme pattern of phenotypic differentiation across subspecies that we observe, with northeastern var. *tuberculatus* ancestry displaying significantly darker purple coloring compared to lighter and greener var. *rudis* stems. This coloration difference among subspecies is consistent with adaptive physiological hypotheses for colder temperatures and northern climates resulting in genetically darker, less reflectant colouring (Chalker-Scott, 1999; Dick *et al*., 2011; Koski & Galloway, 2020). Compared to longitude, fewer latitudinal clines in growth related phenotypes were confounded with genetic ancestry, possibly due to the smaller latitudinal variation sampled, however accounting for ancestry tended to increase latitudinal explanatory power. We found the strongest evidence of latitudinal clines in mid-life history traits—height at flowering, stem width at flowering, and days to flowering (**Table 1; Figure 4**)—a signal of broad-scale geographic adaptation to climate with populations evolved in colder climates growing faster and flowering earlier to avoid severe winters (Stinchcombe *et al*., 2004).

Above and beyond a longitudinal cline in ancestry, we find that var. *rudis* ancestry is preferentially retained (or var. *tuberculatus* ancestry selected against) in agricultural environments, finding on average 8% higher var. *rudis* ancestry in agricultural environments and up 44% more within the most extreme pairing (**Figure 2)**. That the agricultural invasion of *A. tuberculatus* has been more severe in southwestern parts of the range may lead one to predict that var. *rudis* ancestry would be associated with agriculture regardless of a role of selection. However, our paired design—geographically dispersed collections of pairs of natural and agricultural populations <25km apart **(Figure 1)**—explicitly controls for this bias.

Furthermore, that var. *rudis* ancestry is associated with higher male biomass (**Figure 5)** suggests that this ancestry may generally outcompete var. *tuberculatus*. Thus, while controlling for population structure has and continues to be a major challenge for characterizing signals of selection and adaptation (Hoban *et al*., 2016; Barton *et al*., 2019), these processes may not be mutually exclusive, for example in Maize where selective sorting of ancestry underlies adaptation to elevation (Calfee *et al*., 2021). The recent invasion of *A. tuberculatus* into agricultural landscapes over the last century has set up a snapshot of phenotypic and genomic diversity “before and after” (Waselkov *et al*., 2020), providing a rare opportunity for testing not only the phenotypic traits and genomic variation that may contribute to high fitness in newly invaded environments, but the extent to which adaptive variation in agricultural environments may have predated the environmental transition.

We explicitly tested for agricultural pre-adaptation (selective sorting of var. *rudis* ancestry) by examining whether the effect of the proportion of var. *rudis* ancestry on phenotypes varied across treatments, which we designed to mimic key components of natural and agricultural environments. We found that the effect of var. *rudis* ancestry on fitness related traits depended on the reared environment implying locally adaptive subspecies specific variation. However, treatment-specific ancestry effects were not necessarily dominated by var. *rudis* types outperforming var. *tuberculatus* types in the soy competition treatment (i.e. a major axis of agricultural habitats) and underperforming in the water treatment (i.e. a major axis of natural habitats), as we predicted. For male biomass, the ancestry by treatment interaction effect was driven by the strong positive effect of var. *rudis* ancestry on biomass in the control treatment, and relative lack thereof in either the soy or water treatments (**Figure 5)**. Similarly, var. *rudis* ancestry in males led to earlier time to flowering in the control treatment, little difference in the water treatment, and later time to flowering in the soy treatment (**Figure 5)**. One hypothesis for the potential benefit of later flowering in the presence of focal crops like soy or corn is that it may facilitate a longer vegetative growth period allowing *Amaranthus* to dominate the canopy and facilitate efficient pollen dispersal. However, with drought having been shown to select for early flowering genotypes who shorten their life history in response to a shortened growing season (Cohen, 1976; Kozłowski, 1992; Franks *et al*., 2007), our results suggest that the early flowering of var. *rudis* in control treatments—which experienced increased water stress compared to the water treatment—is likely adaptive. Indeed, it is important to note that as opposed to water saturation of the soil, on the hot sunny roof where our experiment was conducted, the water treatment only reduced the severity of soil dry out relative to the control. Thus, the pronounced importance of var. *rudis* ancestry in the control treatment for both flowering-and biomass-based fitness components suggests that var. *rudis* ancestry may experience a selective advantage over var. *tuberculatus* ancestry in drier conditions, as is typical of ruderal habitats, and in agricultural conditions with increasingly frequent droughts. While these hypotheses need further testing in field settings, the over-representation of var. *rudis* ancestry in agricultural environments, higher biomass of var. *rudis* types, and significant environment dependent phenotypic effects of ancestry, lends strong evidence to the role of ancestral pre-adaptation in the *A. tuberculatus* agricultural invasion.

In addition to pre-adaptation, our investigations suggest that on-going local adaptation, but not phenotypic plasticity, is further facilitating any selective advantage that var. *rudis* lineages may have in agronomic environments. We found that regardless of the proportion of var. *rudis* ancestry, natural and agricultural samples showed local adaptation for time to germination, and that biomass was marginally larger and female flowering time marginally later in agricultural environments. The evolution of higher fitness in introduced as opposed to native ranges has often been reported (Leger & Rice, 2003; Erfmeier & Bruelheide, 2005; Caño *et al*., 2008)—consistent with *de novo* adaptation to novel agricultural environments facilitating *A. tuberculatus* reaching a new fitness peak. We find no substantial evidence of increased plasticity in genotypes collected from agricultural habitats compared to those from natural habitats, in contrast to weed generalist “jack of all-trades” hypotheses (Richards *et al*., 2006) that increased plasticity may facilitate the invasion of agricultural environments. On the contrary, we find that var. *rudis* ancestry, which has been preferentially retained in agricultural environments, shows much less plasticity in both biomass and flowering time, facilitating higher fitness in more diverse environments.

This joint inference of the role of ancestry, home environment, reared environment, and geography in shaping patterns of phenotypic variation has thus provided evidence for the invasion of *Amaranthus tuberculatus* into agricultural habitats through local adaptation across multiple timescales.

## CONCLUSIONS

In conclusion, this work has illustrated the power of joint genomic and phenotypic investigation in testing hypotheses about the timescale of adaptation. We find strong evidence for a role of pre-adaptation in the *A. tuberculatus* invasion of agricultural environments, through the preferential sorting of var. *rudis* ancestry, further supplemented by adaptation on more recent timescales. We show that adaptation to agricultural environments has occurred in the face of gene flow, evident by natural-agricultural population proximity predicting similarity of population structure. Future work on the extent of environment-mediated selection for or against gene flow across the genome, the genomic architecture of phenotypic trait differences among environments, and the timescale of allele frequency change associated with agricultural environments will further resolve the enigma of rapid adaptation to human-mediated environmental change.

## ACKNOWLEDGEMENTS

Thanks to James Santangelo, Loren Rieseberg, and the Wright and Stinchcombe labs for feedback on the manuscript. We tremendously appreciate the assistance of the EEB greenhouse staff—Thomas Gludovacz and Bill Cole—at the University of Toronto for the support throughout our experiment, and assistance from many enthusiastic and hardworking undergraduate researchers without which we could have not have performed the common garden experiment, especially Tais Trepanier, Paula Pietraszkiewicz, Xueqian Ma, Eva Gajic, Thomas Lim, and Justin To. A special thanks to Caleb Morse at the McGregor Herbarium, Kansas University for assisting with collections and providing our most westward samples.

Funding for parts of this study was provided by the Society for the Study of Evolutionary Biology Rosemary Grant Advanced Award (JMK), NSERC PGS-D (JMK), Discovery Grants from NSERC Canada (SIW & JRS), and SIW’s Canada Research Chair in Population Genomics.

